# Exploratory Gene Ontology Analysis with Interactive Visualization

**DOI:** 10.1101/436741

**Authors:** Junjie Zhu, Qian Zhao, Eugene Katsevich, Chiara Sabatti

## Abstract

The Gene Ontology (GO) is a central resource for functional-genomics research. Scientists rely on the functional annotations in the GO for hypothesis generation and couple it with high-throughput biological data to enhance interpretation of results. At the same time, the sheer number of concepts (>30,000) and relationships (>70,000) presents a challenge: it can be difficult to draw a comprehensive picture of how certain concepts of interest might relate with the rest of the ontology structure. Here we present new visualization strategies to facilitate the exploration and use of the information in the GO. We rely on novel graphical display and software architecture that allow significant interaction. To illustrate the potential of our strategies, we provide examples from high-throughput genomic analyses, including chromatin immunoprecipitation experiments and genome-wide association studies. The scientist can also use our visualizations to identify gene sets that likely experience coordinated changes in their expression and use them to simulate biologically-grounded single cell RNA sequencing data, or conduct power studies for differential gene expression studies using our built-in pipeline. Our software and documentation are available at http://aegis.stanford.edu.

## Introduction

Since its inception, the Gene Ontology (GO)^1^ has empowered analyses of high-throughput molecular data. Researchers often perform statistical tests using the GO to determine functional enrichments within their discoveries^2–4^ and recently it has also been used to improve model architectures in machine learning applications^5, 6^. The GO hinges on two continuously evolving elements: 1) a collection of curated biological terms with semantic hierarchical relationships and 2) annotations that link genes and gene products to specific terms. When associated with a dataset, such as genes identified from differential gene expression testing^7^, a statistical testing strategy can assign each GO term an “enrichment score”, which measures how genes previously annotated with this term are enriched in the data. The large number of interrelated concepts^8^ and the evolving nature of the knowledge base, while assuring the richness of GO information, also represent a challenge in defining appropriate testing strategies^9^, interpreting and displaying results^10^, and ensuring their reproducibility^11^.

Data visualizations, by illustrating the number of terms, rendering the relations between them and displaying term annotations, can alleviate some of the aforementioned issues. The information in GO is organized as a directed acyclic graph (DAG), where each node corresponds to a GO term and edges indicate relations such as “part of”, “a type of”, etc. It is then natural to rely on the DAG structure of the GO for information visualization (Supplementary Figure 1). Indeed, there are a number of published tools that leverage the structure and can be broadly divided in two groups: ones that are specialized for local graph characteristics, and ones that are optimized for rendering global graphs structures.

On the one hand, many existing GO tools focus on displaying a small subset of terms to reveal nuances of the hierarchical relationships. For instance, one of the most commonly-used web interfaces, QuickGO^12^ allows querying and browsing a single GO term along with its related terms (and annotations) as a DAG. REVIGO^13^ can present multiple selected terms in a semantic graph via different representations (e.g., its *scatter and table* view, *tree map* view, and *interactive graph* view). Other tools feature displays where node and links can be retrieved with more interactive features^14–16^, or where node enrichment scores can be highlighted with additional customization^17–19^.

On the other hand, relatively fewer visualizations aim to simultaneously displaying the entire GO^20, 21^, despite extensive literature on large graph or network visualization^22–26^. In contrast to small graph displays, these tools are typically not flexible enough to highlight node- or link-specific details due to numerous visual elements. However, they can provide a global view of the graph, such as nodes clusters or overall hierarchies that are especially helpful for understanding trends of term relationships or enrichment scores in the GO.

Our new open-source software AEGIS (Augmented Exploration of the GO with Interactive Simulations) aims to bridge the merits of both visual approaches, and support extensive interactions with the information coded in the ontology. We link representations of local and global structures within the GO DAG by adopting the *focus-and-context* framework, reminiscent of classical principles in visual information system design: *overview first, zoom and filter, then details-on-demand*^27^. AEGIS capitalizes on the flexibility and power of efficient search routines to render dynamic displays for real-time interactions. Not only can the users interact with the interface to visualize their results output from existing pipelines with increasing degrees of specificity, they can also choose any vantage point to explore the GO with our visualization prior to collecting their data or running their pipelines. During the exploratory process, AEGIS allows them to extract biological information relevant for simulations and hypothesis generation, as well as power calculations for study design.

## Results

To increase interpretability and facilitate user interactions with the GO, we reasoned that it is key to both effectively display granular information relative to a set of concepts of interest and provide a means to put them in the larger context of the ontology. AEGIS achieves this objective by a new visualization: *focus and context graphs*. It couples a Sugiyama-style graph^28^ (*the focus graph*) that renders the hierarchical structure of a selected sub DAG with a silhouette view (*the context graph*) that provides indication of the overall number of nodes that are similar to or related to the concepts in the sub DAG. Both the focus and context graphs are customizable in real time thanks to our customized data structure and software system (Methods and Supplementary Figures 2-4). AEGIS complements the focus and context graphs with interactive highlighting options and layouts, including the *buoyant layout*, a graph drawing strategy that incorporates gene annotation information. The buoyant layout relies on a novel algorithm we developed, and improves the interpretation of the hierarchical levels in the GO terms that are assigned to the same level share a similar number of annotated genes (Methods, Supplementary Algorithms, and Supplementary Note 1).

Unlike most GO visualizations that are used at the last step of the analysis to illustrate results, our visualization strategies allow researchers to explore the ontology and plan experiments prior to data acquisition. With interactive features complementing the focus and context graphs, they can generate biologically plausible simulated data and run power studies: in fact, interaction in AEGIS is not limited to customization of graph displays, but includes multiple methods to identify a set of concepts of interest, generate hypotheses, and select study parameters (Methods and Supplementary Note 2). Unique to AEGIS, the context graph can conveniently summarize the multiplicity of the tested hypotheses within the hierarchy, and the focus graph can magnify precise positions of the non-null hypotheses which researchers aim to discover. Subsequently, they can visualize the dependencies among the hypotheses and identify potential testing biases. As new testing strategies are emerging^29, 30^, these insights are valuable for benchmarking their performance and comparing their underlying assumptions.

We next resort to a number of biological examples that showcase the different aspects of our visualization and its applications. While the main results below are renderings of static displays, we refer the reader to detailed video tutorials and documentation of the examples (Supplementary Videos and Supplementary Note 3), as well as a lite on-line interactive platform available at http://aegis-viz.appspot.com.

### Focus and context graphs

The focus and context graphs are two juxtaposed representations tailored to balance the global structures and local details of a DAG with tens of thousands of nodes and links. The focus graph displays precise term relationships and annotation information via a *bona fide* graph representation following the Sugiyama style: nodes are arranged in different levels, with the roots at the top and a parent always above its children so that all links are directed downwards (Figure 1, left hand side). We will discuss in the next section the different options for assigning a node to a level, but it is important here to remark that levels are at the basis of the coupling of the focus and context displays. The context graph is represented by a bar chart (Figure 1, right hand side), capturing the node counts at each level of a larger reference DAG (by default the entire GO). The coupled displays allow the viewer to understand how many nodes are on the same level of a concept of interest, and how many are at higher or lower levels in the hierarchy.

**Figure 1:**
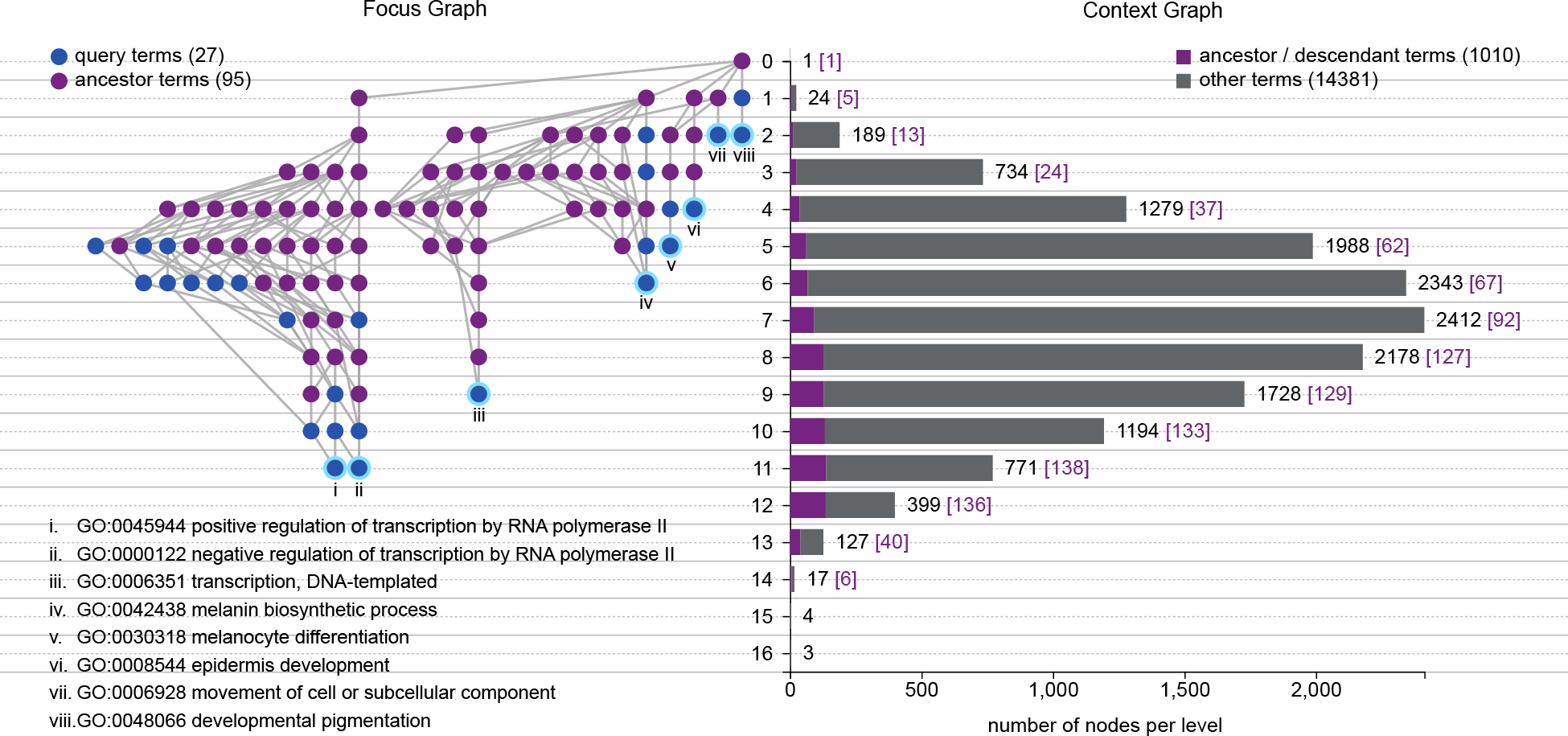
Focus and Context Graph Representations of AEGIS. The GO DAGs are rendered by focus and context graphs in AEGIS based on specific term queries. In this example, the focus graph on the left includes a total of 27 significantly enriched functions, which are highlighted in blue; all of their ancestral nodes under “biological processes” are displayed in purple. The least redundant leaf terms are identified with roman numerals, and annotated with their GO identifiers. We use the root-bound layout, where the level of a node corresponds to its longest distance to the root. The position of each node within its level in the focus graph depends on semantic relationships: related nodes are grouped together. The context graph on the right captures the entire GO DAG under the root of “biological processes” with 15391 terms; the silhouette view indicates the number of nodes in each level under the root-bound layout. Among the nodes in each level, the count of the terms that are ancestral or descendent to the 27 terms (including themselves) are highlighted in purple.

To provide a concrete illustration, we used AEGIS to create a visualization of 27 significantly enriched GO terms for hair color phenotypes from a genome-wide association study^31^ (Supplementary Note 3). By displaying the ancestral terms, it becomes apparent that some of these concepts are more closely related than others: the focus graph on the left hand side of Figure 1 identifies with roman numerals the most specific of the 27 terms, and groups together the concepts with a most recent common ancestor. Aside from stylistic variations and possible customizations, the focus graph resembles standard representations of the GO DAG^3, 12, 28^. The context graph on the right hand side, in contrast, is unique to AEGIS: the silhouette representation of the entire “biological process” ontology DAG (15,000 terms) allows us to indicate the locations of the approximately 1,000 concepts related to the 27 query terms, highlighting, for example, how they include 907 descendants.

The focus and context graphs are achieved without overcrowding the display with numerous objects and without appreciably increasing the computing time. Consequently, the gains in computational and representational efficiency facilitate real-time interactions with the GO DAG. For example, an investigator can explore focus graphs anchored on different terms, or customize selections of context graphs (Methods and Supplementary Videos).

### Buoyant layout that incorporates annotation

The visualization in Figure 1 exemplifies a standard option for assigning a node to a level in a Sugiyama-style graph^28^: the level represents a node’s longest distance to the roots. We refer to this leveled layout as the *root-bound* layout. Another standard option assigns a node to the longest distance to the leaves—we call this the *leaf-bound* layout, and both options are available in AEGIS. The *root-bound* and *leaf-bound* layouts achieve spatial compactness by minimizing the number of levels, but also have their limitations. A viewer is naturally prone to interpret terms in the same level as sharing the same degree of specificity. This might be approximately true in some contexts, but it is rather misleading in the case of the GO DAG.

The ontology is created as a way of representing existing literature and the curation process focuses on term-to-term relationships, not the overall graph structure. While it is always true that descendant terms are more specific than their ancestors, the number of terms separating a concept from the root or leaf nodes is not a reliable indication of its specificity. Some biological processes have been studied in detail, so that a number of different terms exist to describe them, while others are known at a much less granular level. One can imagine a well studied concept separated from the root node by 10 concepts, while a less studied concept of comparable specificity might be only two nodes away from the root.

Fortunately, in the case of GO, we have another source of information on the level of specificity of a node: the number of genes that annotate it. AEGIS leverages this in a novel Sugiyama-style graph, the *buoyant layout*, which enjoys two properties: (1) a parent node is always placed at a level above its children; and (2) a node with fewer annotations is never placed higher than any node with more annotations (Methods and Supplementary Note 1).

To contrast the renderings of the different layouts, we re-analyzed a chromatin immunoprecipitation sequencing (ChIP-seq) study^3, 32^ (Supplementary Note 3), and selected two significant terms to highlight their differences (Figure 2). In the root-bound layout, for instance, the term “ruffle” (with 164 annotations) is at a higher level, and therefore appears to be more generic than the term “actin cytoskeleton” (with 448 annotations) only because a smaller number of concepts lie between “ruffle” and the root “cellular component”. The buoyant layout, in contrast, preserves a more interpretable node ordering based on the number of annotated genes. We also remark that the buoyant layout is particularly meaningful when coupled with the context graph: each level corresponds to a range of gene annotation counts (Supplementary Videos).

**Figure 2:**
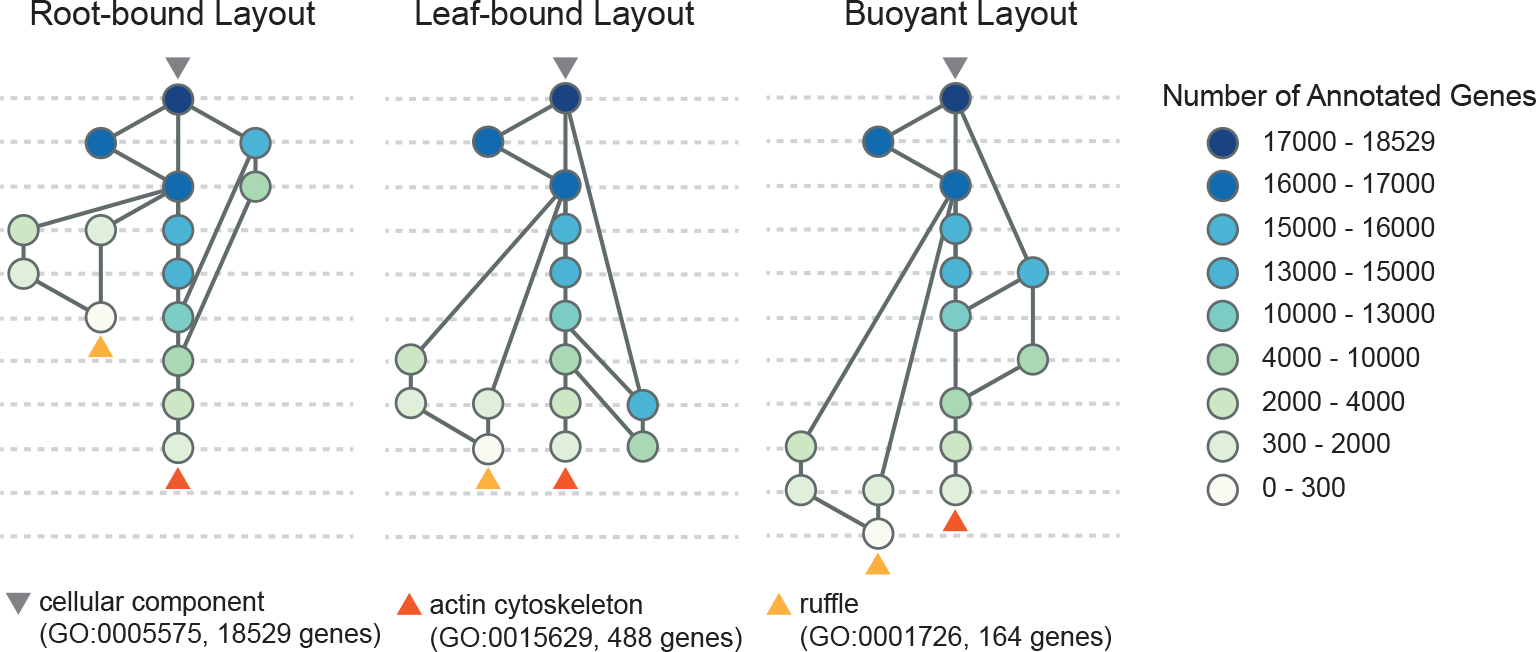
The Buoyant Graph Layout for the GO. Root-bound, leaf-bound and buoyant layouts. Each link is assumed to point downwards from the root (indicated with a gray triangle), and the nodes are colored depending on the number of genes with which they are annotated (or the node size), as in the legend on the right hand side. Orange and yellow triangles point to two significant cell components identified from the CHiP-seq study by Valouev et al. Note that the root-bound layout places ruffle at a higher level that actin cytoskeleton, despite the fact that ruffle is annotated with 164 genes, and likely a more specific compartment than actin cytoskeleton, which is annotated with 488 genes. Similar size-level inconsistencies appear in the leaf-bound layout, but the buoyant layout successfully resolves these issues across all the nodes.

### Interactive features and Gene Ontology navigation

By choosing which nodes to display in the focus graphs, against which context to interpret them, the user can navigate the GO and extract annotation information useful for simulations and study designs. This process is further enhanced with many auxiliary interactive features (Methods and Supplementary Videos). Here we start with showing how AEGIS facilitates extracting information from the GO for the purpose of simulations. We consider the task of generating artificial signals from single cell RNA sequencing (scRNA-seq), where one is interested in discovering subpopulations of cells that have distinct expression signatures. Rather than arbitrarily deciding which genes differ across simulated cell types^33, 34^, AEGIS allows a researcher to leverage the GO to identify a group of genes that are biologically related, specifying a signal structure that is realistic, and model a cellular process the researcher is specifically interested in.

We simulated the transcriptomic profiles of single cells that differ by cell cycle states, a common confounding factor in scRNA-seq data (Figure 3). Previously, cell cycle genes have been collectively considered at the resolution of more generic GO terms such as the cell cycle process (GO:0022402)^35^. With AEGIS we can explore and select more detailed sub-processes: specifically we identified three descendants: G1/S transition of mitotic cell cycle (GO:0000082), G2/M transition of mitotic cell cycle (GO:0000086), and mitotic cell cycle arrest (GO:0071850). We select then 10 genes among the ones annotating each of these terms and refer to these 30 genes as the “signature genes”. Using standard procedures (Supplementary Note 3), we can then generate expression data for 120, 150, 100, 300 cells for G1/S transition, G2/M transition, cell cycle arrest, and control respectively. Each cell type has higher expression of its 10 signature genes, while the baseline cell type does not express any of these transition markers.

**Figure 3:**
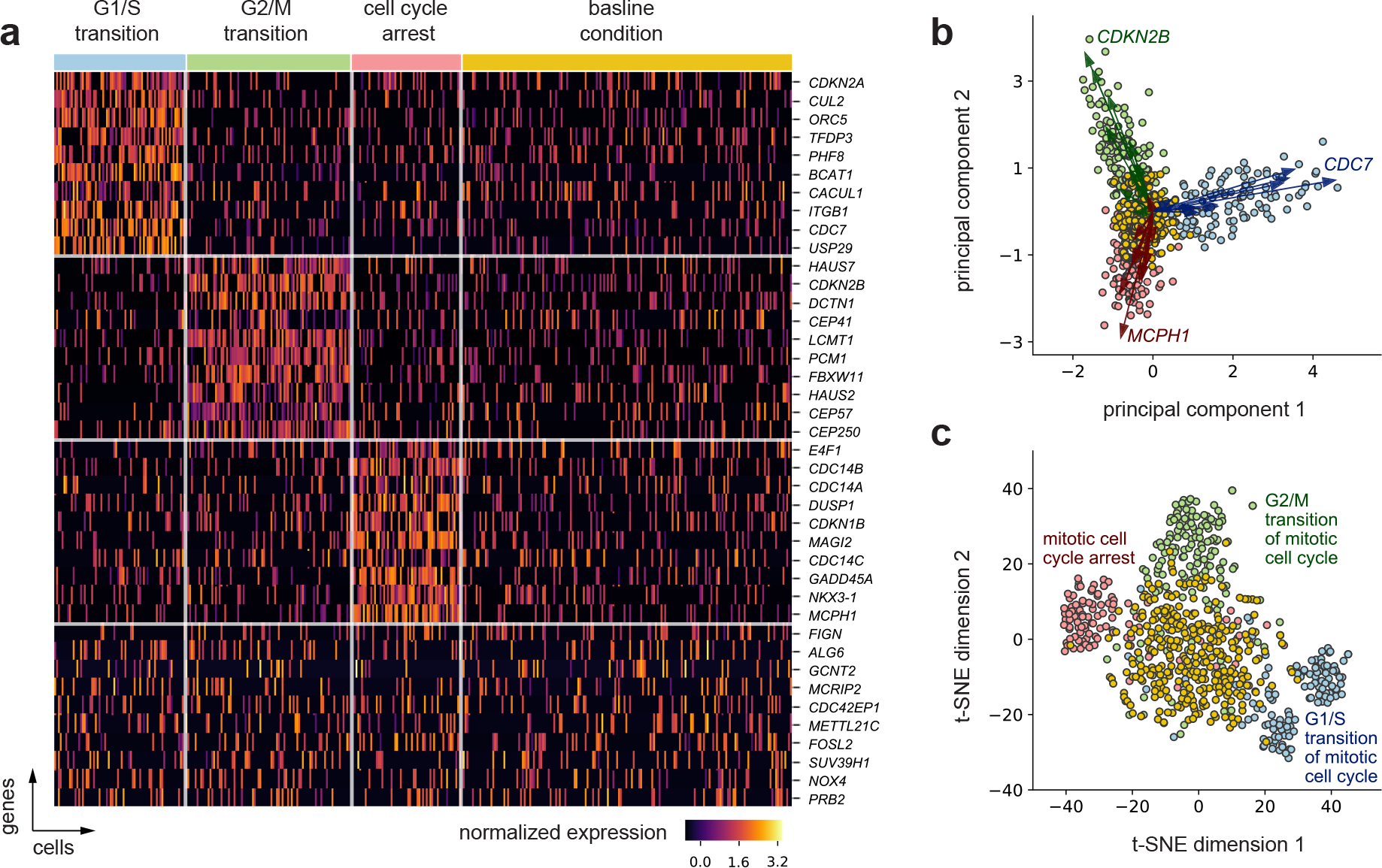
Simulated scRNA-seq with Cell Cycle Signatures. (**a**) Heatmap. Gene expression signatures for cells in G1/S transition (120 cells), G2/M transition (150 cells), cell cycle arrest (100 cells), and a baseline cell type (300 cells, with no cell cycle signature). The first 30 genes are cell type specific signatures used for the simulation: 10 randomly selected from each of the set of gene annotating the three GO terms corresponding to G1/S transition, G2/M transition, and cell cycle arrest. The last 10 genes are selected at random from the total of remaining 2030 genes for which expression was generated using a zero-inflated negative binomial models with mean parameters varying with cell types. (**b**) Biplot on the top 2 principal components (PCs). The biplot includes the first and second PCs and each point represents individual cells projected in this space. The vector arrows are the projection of the 30 signature genes colored by their annotation group, and those with largest magnitude per group are indicated. The point coordinates and the gene vectors were scaled (while preserving the inner products) to fit the display space. (**c**) t-SNE 2D embedding. The t-SNE visualization of the cells also highlights the four distinct cell sub-populations and processes they are enriched in. The t-SNE was computed based on the top 10 PCs of the data. The scale of the t-SNE plot is arbitrary.

A biologically sound ground truth can help contrast alternative visualizations of the data, such as PCA^36^ and t-SNE^37^ (Figure 3). As we expect, both methods organize the data points in a way consistent with the ground truth cell types, with better cell type segregation under t-SNE. However, our functionally related gene set uncovers how visualizations with PCA can offer additional insights: PCA-based biplots can indicate directions in the data dominated by individual genes. These “gene directions” can in turn suggest “functional directions”, i.e., annotated gene sets that collectively align in the subspaces spanned by the top principal components.

### Power analysis for GO enrichment testing

Study design is one of the contexts where it is most useful for an investigator to interact with GO visualizations and AEGIS further enables them with a built-in power simulator, filling in a surprising gap in the literature of enrichment tests^2, 18, 38, 39^ (Methods). While enrichment hypotheses are stated and tested at the level of GO terms, their truth status depends on the expression of the genes associated to the concepts: because often the same gene contributes to the annotation of multiple GO terms, there are logical dependencies between enrichment hypotheses. Without actual access to the GO structure, it is difficult to simulate these dependencies, which have important consequences for power and interpretability of results.

AEGIS’ pipeline aids the investigator in multiple ways (Supplementary Figure 5): 1) By allowing navigation of the GO, it enables the investigator to select ontology terms that are representative of the biological process under study and determine that all or some of the genes in their annotation are differentially expressed—thereby anchoring the power study to biologically relevant hypotheses. 2) Once differentially expressed genes are specified, AEGIS identifies all the non null enrichment hypotheses, according to one of two standard definitions (e.g., ‘competitive’ or ‘self-contained’^9^, described in Supplementary Note 2), allowing the investigator to evaluate the specificity of the hypotheses tested. 3) The built-in power simulator generates gene expression datasets using the sample size and signal strength specified, carries out testing with a choice of testing and multiplicity correction strategies, and computes power and False Discovery Rate (FDR). 4) The power results are displayed with reference to the relevant GO sub-DAG, using *binder plots*. By organizing nodes linearly while still displaying hierarchical information, this visualization facilitates the display of additional node-specific information such as term names, bar plots or heatmaps for the summary statistics from the power analysis. The order of nodes is based on number of annotations (following the bouyant layout), and edges are rendered with curves that preserve Sugiyama-style graph drawing rules.

As an example, we conducted a power analysis to guide a study with the goal of confirming the enrichment of heart-specific developmental processes (Supplementary Note 3). We study how power changes with sample size using two different strategies for formulating and testing the enrichment hypotheses: competitive nulls with hypergeometric test, and self-contained nulls with p-values obtained with Simes’ combination rule (Figure 4 and Supplementary Note 2).

**Figure 4:**
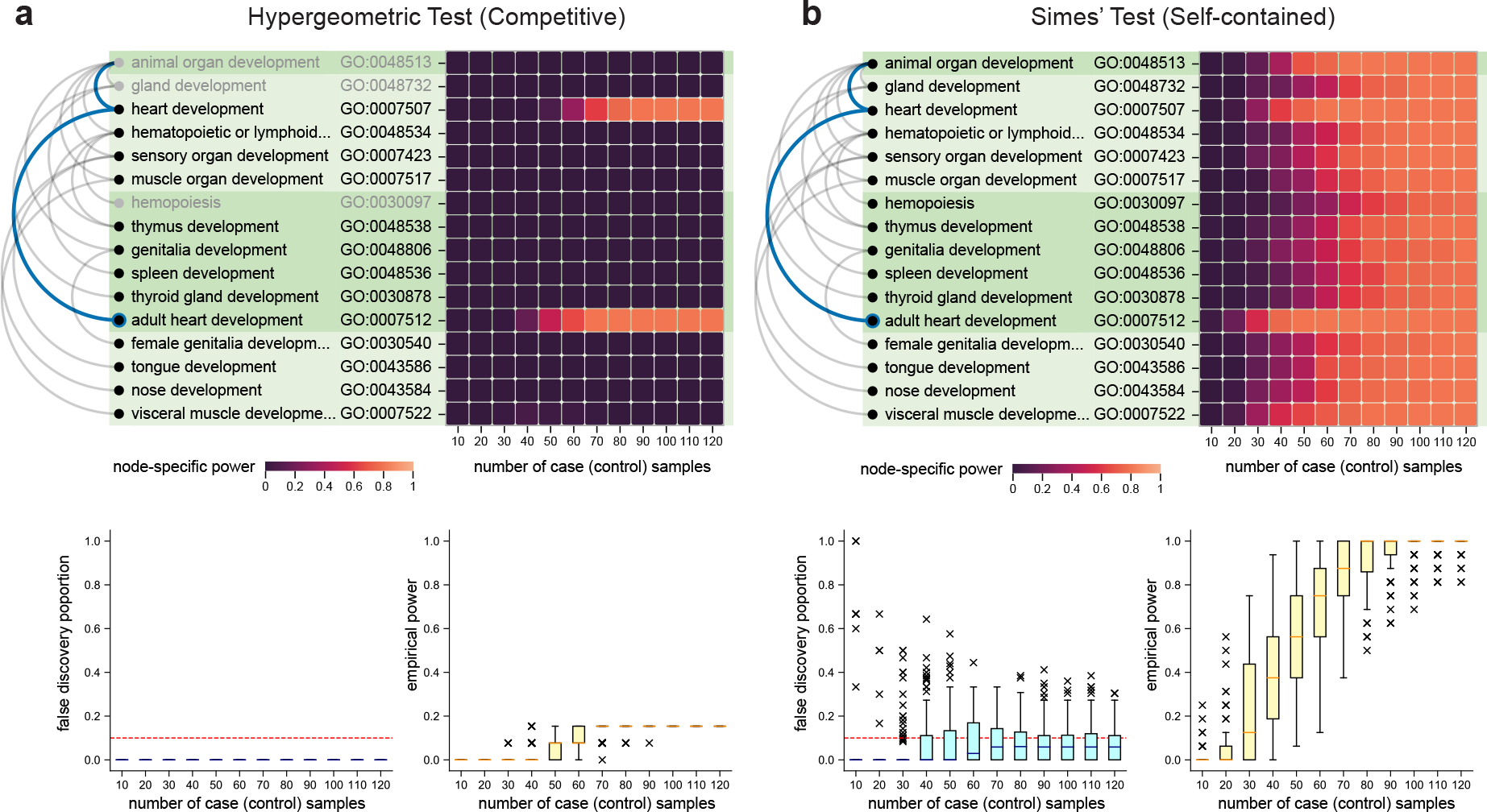
Binder plots, Heatmaps and Box Plots for the Heart Development Case Study. (**a-b**) All the genes from those annotating the term “adult heart development” (with node border marked blue and ancestors indicated in blue as well) and defined as differentially expressed. This defined a set of non null hypotheses for the competitive and the self-contained framework, all displayed in the binder plots, rooted at “animal organ development”. In each binder plot, the non-null terms relative to the appropriate hypothesis are shown with black text: for completeness and ease of comparison, binder plot for the competitive tests includes in gray three null terms related to the non-nulls. The terms are ordered by descending node sizes, and the different levels are distinguished by alternating shades of green. The heatmaps highlight the fraction of times a particular node is rejected across 100 repetitions per sample size regime. (c) The box plots in the bottom portion of the displays summarize the false discovery proportion (at level 0.1) and empirical power across all nodes. The Benjamini-Hochberg procedure was used for multiple testing adjustment.

As previously documented^9^, increasing the number of samples leads to monotonically higher power for self-contained tests, but can have limited impact on competitive tests that are based on a “gene sampling” (rather than subject sampling) strategy. On the other hand, inspection of the binder plots reveals that the loss of power of competitive/hypergeometric tests is mostly due to their not rejecting rather non-specific hypotheses: they do single out “heart development” (where the signal was planted) and “adult heart development” (whose annotation is very similar).

We note that this strategy can be used to benchmark multiplicity-correction procedures, and documented the results of a comparison of DAGGER^29^, a recent sequential testing method, and the Benjamini-Hochberg procedure (Supplementary Note 3).

## Conclusion

AEGIS employs multiple strategies to navigate the GO and present GO-based discoveries: the focus-and-context graph, the buoyant layout, and the binder plot are novel visualizations that lever-age graph-theoretic properties of the GO DAG, extensive customization and interactive displays allow user engagement, and built in power simulations facilitate study design.

AEGIS is available both as a web-based application for simple GO exploration, and a downloadable software package for hypothesis testing and further customization. Currently, version-controlled visualizations rendered via AEGIS can be downloaded as vector graphics, such that attributes such as colors, annotations and node positions can be customized via common editing tools. The computation back-end and visualization front-end, along with online documentations, tutorials and videos, are open-source and portable. As such, existing GO analysis pipelines can employ the functionalities of AEGIS to present and disseminate their results. For example, our current pipeline is compatible with emerging GO testing strategies and multiple comparison procedures^3, 29, 30^, and it also has the flexibility to support the analyses of a broader range of data input types with these options. Moreover, the visualization strategies can potentially be transferred to other ontologies^3^, such as the Plant Ontology and Human Disease Ontology. For ontologies where annotations are not limited to gene sets, we can extend the buoyant layout to encode other side information associated with each term to render interpretable levels graph layouts using their specific constraints.

## Methods

### 1 GO data preprocessing

The analysis in this manuscript is based on the GO version: release-2018-06-21 (format version: 1.2) with gene annotations downloaded from the National Center for Biotechnology Information (NCBI) gene2go table (ftp://ftp.ncbi.nih.gov/gene/DATA/gene2go.gz, release-2018-06-21). The open-source software goatools (https://github.com/tanghaibao/goatools) was used to parse these files. For the associations, all evidence types are included as default^3^. All relationships (e.g., “is_a”, “part_of”, “has_part”, or “regulates”) for the GO terms were used for the cellular component ontology, but only “is_a” relationships were kept for the biological process ontology.

To handle the redundancy of GO terms during hypothesis testing, we developed a *DAG refinement* algorithm, which also preserves the semantic relationships in the original DAG (Supplementary Algorithm 1). Essentially, the algorithm iteratively remove a parent if one of its child is annotated to the exact same gene set. DAG refinement is optional for visualization but is strongly recommended for hypothesis testing, because if a parent and child share the exact same gene sets, the corresponding test statistics would be identical and unnecessarily increase the burden of testing multiplicity. Further, the user can optionally filter the node-size range to remove overly generic or specific terms, a function commonly used in gene set testing tools^3^. We found that other works rely on DAG reduction approaches for similar reasons^3, 6^, however, we were unable to identify a full algorithmic description or justification of the process. Thus, our algorithmic description is also accompanied with its theoretical justification (Supplementary Note 1).

### 2 Customization of the context graph

The context graph is by default the entire GO, but it can be narrowed to a sub-DAG by specifying a set of *context anchors*: any GO terms can be chosen as anchors, playing one of three roles in identifying the sub-DAG: *root, leaf* or *waypoint*. Selecting root anchors will result in a DAG that includes all their descendants; leaf anchors in a DAG that only include their ancestors; and waypoint anchors in a DAG which include both their descendants and ancestors (Supplementary Figure 2). This procedure can be seen as a way to eliminate nodes that have no semantic relationship with the context anchors. The induced sub-DAG can be further processed using DAG refinement.

Once the context is specified, the focus graph can be similarly specified with *focus anchors*, which can be defined as *waypoint* or *leaf*. If these anchors are not specified, the default would be the the context anchors as leaf anchors. In order to make sure that the number of nodes in the focus graph is small enough to allow interaction, two constraints are implemented on the sizes of the displayed sub-DAG: 1) we limit the number of focus anchors; and 2) descendants beyond the immediate children of the focus anchors are displayed only when the the size of the resulting focus graph is predicted to be smaller than a threshold. Both these parameters can be adjusted.

AEGIS supports three layouts for the focus graph: *root-bound*, *leaf-bound*, or *buoyant*. The first two are standard topological layouts^28^, while the *buoyant* layout is new and described below. Given a context graph, one can change the focus anchors in the focus graph in multiple interactive ways, including key word search, node selection, or random draws of nodes from a specific level (Supplementary Videos).

### 3 Buoyant layout and the bubble float algorithm

For graphical displays of the GO DAG, a buoyant layout requires two constraints: each parent node should be placed at a higher level than all of its children (*topological constraint*) and a node annotated with a smaller number of genes should never be placed higher than any larger node (*descending node size constraint*). The *bubble float algorithm* aims to generate the buoyant layout based on the two constraints. It starts by initializing the layout by lexicographically sorting all the nodes according to (1) descending ordering of the node size and (2) ascending ordering of the longest distance to the root. The remainder of the algorithm consists in repeated applications, starting from the bottom of the graph, of a *float* operation that merges multiple nodes into a layer so long as the two constraints remain satisfied. We show that the *bubble float algorithm* satisfies both the topological constraint and the descending node size constraint during initialization and at each iteration. In addition, the necessary and sufficient conditions of the buoyant layout are proven theoretically, such that it is guaranteed to exist for any GO DAG (Supplementary Note 1).

The resulting number of levels of the buoyant layout is always equal to or greater than that of the root-bound layout (Figure 2), because the latter only requires the topological constraint. Particular to the GO, this can be advantageous when too many nodes are displayed and could be more spread out by levels. Thus, we also provide a parameter to control the maximum span of node sizes on each level (Supplementary Algorithm 2).

Once the level of each node in the DAG is determined, what remains is the ordering of the nodes in each layer for the focus graph. Computing the optimal ordering of nodes in each level that minimizes edge crossings between layers reduces to the problem of *bipartite crossing minimization*, which is unfortunately NP-hard^40^. Yet, there exist many graph drawing heuristics, which we build upon to determine the layout of the focus graph: we group anchors together if they are semantically related, add exclusive ancestor and descendant nodes to the respective groups, and separate these groups the nodes during display (Figure 1, Supplementary Algorithm 3). The grouping delivers a clearer interpretation of the focus anchors and their relatives. In contrast to the focus graph layout, the rendering of the bar layout of the context graph does not require node ordering at each layer. Because the representation only concerns the node counts, the computation cost for the context graph is negligible.

### 4 Software architecture

The specifications of the context and focus graphs, choices of different layouts, and states in the interaction process can be combinatorial. To efficiently support the graphical rendering and interaction, we created a new data hierarchy: a data object includes 1) primary attributes that are invariant across any context and focus representations; 2) secondary attributes that can vary under different choices of the context graph, but are invariant to what the focus graph is; 3) tertiary attributes that can vary with the choice of the focus graph and interactions on the graph. All these attributes collectively define how an object is displayed on the front-end interface, but because some of the attributes can be expensive to compute, we use this hierarchy to reduce the space and time needed for computations (Supplementary Figure 3). It is expected that certain computations in back-end, such as performing a statistical testing method can be more computationally expensive. Nevertheless, this computation does not affect the layout—our hierarchy assigns them to primary attributes, and none of these results need to be recomputed for layout and interaction purposes. Therefore, we were able pre-compute and cache the simulation results prior to running the web server. Another computational bottleneck in our system is data communication with the GO database and basic file parsing (which can take up to minutes depending on Internet connection).

To systematically integrate this data hierarchy, we created new software libraries to define the data classes and perform the core algorithms (Supplementary Figure 4). We also utilized state-of-the-art Python packages to query and parse GO ontology databases, and incorporated Flask, a Python micro-framework to deliver server side computations. All of the user interactions were developed based on commonly-used JavaScript packages: jQuery.js and D3.js to support lightweight front-end computations on the client side.

Despite the size of a DAG with tens of thousands of nodes, our computational framework and system architecture offer the advantage of minimal time and space required for interactions with the full-scale data. As a result, our application does not require expensive resources to perform all the computations in the background, and reduces the need to be hosted as a centralized web portal. All the development and demonstrations were performed on a MacBook Pro laptop with a 2.8 GHz i7 processor with 8 Virtual Cores and 16GB RAM. The interaction updates in the front-end interface each take less than 0.1 seconds.

### 5 Power analysis workflow

The power analysis for GO enrichment testing in RNA-seq studies requires three ingredients: 1) the ground truth: *a priori* defined truly differentially expressed genes, as well as whether each GO term is a true null or non-null hypothesis; 2) the data: the gene expression matrix and term-specific p-values; and (3) the statistical procedure: testing methods to select which terms are significant. The selected ground truth genes determine if a GO term is a true null hypothesis or not, and there are two distinct types of null hypothesis to be considered: the *self-contained* null and the *competitive* null^9^. We say that a GO term is a self-contained null if no gene associated with the term is truly differentially expressed; a term is a competitive null if the genes associated with this term are at most as often differentially expressed as the genes not associated with this term (Supplementary Note 2).

To generate the data, AEGIS takes the selected truly differentially expressed genes, effect size *β*_effect_ and sample sizes *n*_control_, *n*_case_ as parameters to generate *g*-dimensional gene expression matrices for the control samples and case samples. For simplicity, the gene expression values for the controls are sampled from 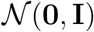, a multivariate normal distribution with a length-*g* zero mean vector and a covariance matrix equal to the identity matrix; and the cases are similarly sampled from 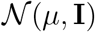, where *μ* is vector of values equal to *β*_effect_ only at the truly differentially expressed genes and 0 otherwise. This vanilla model can be extended for more complicated scenarios: for example, the single-cell RNA-seq simulation uses the negative binomial model instead and impose additional zero values in the expression matrix, also known as as “drop-outs” (Supplementary Note 3). Further, the assumption that the gene measurements are uncorrelated can be relaxed given the knowledge of platform-specific noise patterns.

For testing, a total of *g* gene p-values are obtained based on a two sample t-test; these are the ingredients to determine GO term-specific p-values. For the self-contained hypothesis, the Simes’ combination rule aggregates all the gene p-values annotated with a GO term to obtain the term p-value; For the competitive hypothesis, the hypergeometric test is based on first selecting significant genes based on a gene-wide significant threshold, and testing whether these genes are independent of those annotated to a specific GO term (Supplementary Note 2). For either case, we control the false discovery rate at level *q* using the Benjamini-Hochberg (BH) procedure. Note that this pipeline is modifiable: other methods, such as Bonferroni correction or DAGGER, can replace BH for multiplicity correction (Supplementary Note 3).

Without loss of generality, let 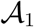 be the set of non-null terms for a particular type of null hypothesis, and let 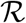 be the set of terms rejected by the appropriate procedure. Then, the (empirical) power and false discovery proportion (FDP) are defined as follows:

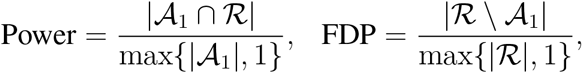

where |·| denotes the size of the set, and \ denotes set difference. For a particular power calculation, we fix the ground truth and repeat the data generation process to obtain multiple realizations of Power and FDP, in order to infer the power as well as the false discovery rate (FDR) of detecting the true non-nulls.

